# Structural changes in perineuronal nets and their perforating GABAergic synapses precede motor coordination recovery post stroke

**DOI:** 10.1101/2022.10.05.510951

**Authors:** Egor Dzyubenko, Katrin I. Willig, Dongpei Yin, Maryam Sardari, Erdin Tokmak, Patrick Labus, Ben Schmermund, Dirk M. Hermann

**Author notes:** Corresponding authors Egor Dzyubenko,; Dirk M Hermann.

## Abstract

Stroke remains one of the leading causes of long-term disability worldwide, and the development of effective restorative therapies is hindered by an incomplete understanding of intrinsic brain recovery mechanisms. Here we explored how perineuronal nets (PNNs), the facet-like extracellular matrix layers surrounding fast-spiking interneurons, contribute to neurological recovery after focal cerebral ischemia in mice with and without induced stroke tolerance. Due to the insufficient resolution of conventional microscopy methods, the contribution of structural changes in PNNs to post stroke brain plasticity remained unknown. Using superresolution stimulated emission depletion (STED) and structured illumination (SR-SIM) microscopy, we revealed that PNN facets become larger and less dense in the post-acute stroke phase. These morphological alterations in PNNs are transient and correlate with the increased surface of contact between activated microglia and PNN-coated neurons. The transient loosening of PNNs after stroke allows for dynamic reorganization of GABAergic axonal terminals on inhibitory interneurons in the motor cortical layer 5. The coherent remodeling of PNNs and their perforating inhibitory synapses precedes the recovery of motor coordination after stroke and depends on the severity of the ischemic injury. Our data suggest a novel mechanism of motor cortical plasticity after stroke, and we propose that prolonging PNN loosening during the post-acute period can extend the opening neuroplasticity window into the chronic stroke phase.

**Highlights:**

- PNNs are degraded partially and transiently post-stroke
- Transient attenuation of PNNs correlates with GABAergic synapse remodeling
- Transient attenuation of PNNs precedes functional recovery post stroke
- Activated microglia preferentially contact PNN-coated neurons post stroke

**Graphical Abstract:** 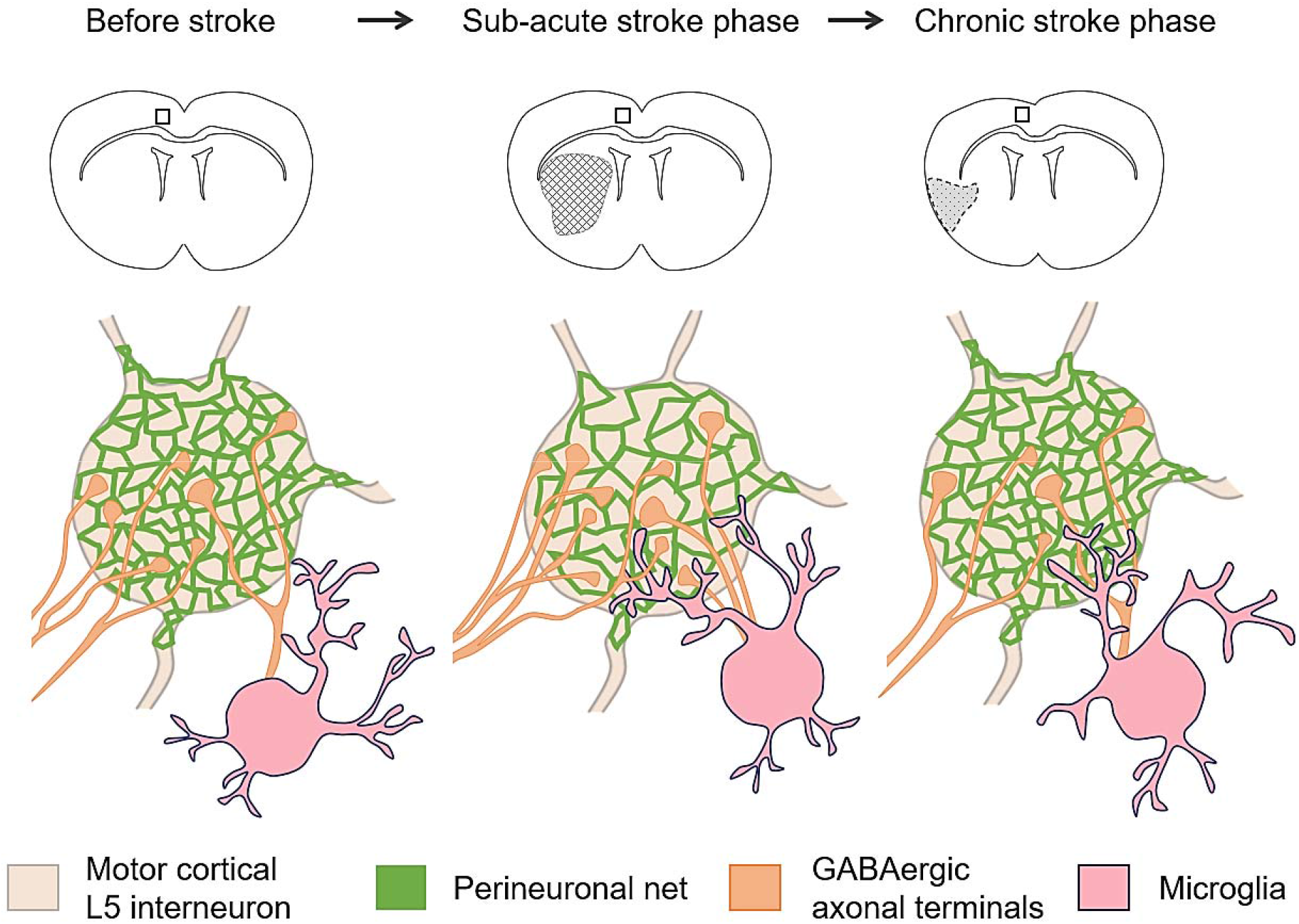

## Introduction

Brain remodeling is essential for regaining compromised motor activity and coordination post stroke. Neurological recovery after stroke involves several neuroplasticity mechanisms including corticospinal tract rewiring (Hermann and Chopp, 2012), sprouting of interhemispheric cortico-cortical projections (Liu et al., 2010), and remodeling of local intracortical connectivity (Carmichael et al., 2001). Motor cortical activity defining motor commands during skilled limb movements is selectively distributed across neurons with distinct projection patterns (Park et al., 2022), suggesting that reorganization of both long-range and local connectivity is similarly pivotal for a successful recovery after stroke. Experimental treatments promoting pyramidal tract plasticity have been proposed (Reitmeir et al., 2011), but harnessing the plasticity of intracortical projections requires a deeper understanding of local connectivity changes in the motor cortex post stroke. Cortical oscillations underlying motor learning arise from the fast-spiking activity of layer 5 (L5) interneurons (Otsuka and Kawaguchi, 2021), which are the main source of inhibition in neocortical microcircuits (Naka and Adesnik, 2016; Packer and Yuste, 2011). Although motor cortical L5 interneurons are critical for controlling coordinated movements, their involvement in post stroke brain remodeling has not been systematically studied to the best of our knowledge.

Cortical L5 fast-spiking interneurons express parvalbumin (PV), and potassium channels with rapid activation and deactivation kinetics (Kv3.1) and are coated with perineuronal nets (PNNs) on the extracellular side (Chow et al., 1999; Hartig et al., 1999; Matsuda et al., 2021). PNNs are condensed lattice-like layers of extracellular matrix (ECM) composed of multiple macromolecules including hyaluronic acid, chondroitin sulfate proteoglycans (CSPGs), and link proteins (Carulli et al., 2006; Su et al., 2019). These polymeric assemblies propagate to the extracellular space (ECS) and are anchored to the neuronal surface via hyaluronic acid synthases (Frischknecht and Seidenbecher, 2008; Kwok et al., 2010). PNNs are rigid structures resistant to chemical decomposition (Deepa et al., 2006) that are formed in an activity-dependent manner (Dityatev et al., 2007) and restrict synaptic plasticity (Frischknecht et al., 2009; Pizzorusso et al., 2002). They compartmentalize neuronal surface, limit astrocyte-neuron direct membrane contacts and new synapse formation, stabilize existing connectivity, and hypothetically contribute to potassium buffering (for review see (Dzyubenko et al., 2016; Fawcett et al., 2019)). Thereby, PNNs maintain the fast-spiking properties of interneurons (Tewari et al., 2018) and inhibitory control in neuronal networks (Dzyubenko et al., 2021).

In the adult brain, CSPGs within PNNs restrict neuronal plasticity by inhibiting axonal sprouting (Galtrey and Fawcett, 2007), and experimental approaches involving non-specific ECM digestion have been shown to promote functional recovery post stroke (Gherardini et al., 2015). It is unlikely though that the complete removal of ECM and PNNs in particular can be implemented clinically because it induces epileptiform activity (Arranz et al., 2014; Balashova et al., 2019) and impairs memory formation (Gogolla et al., 2009; Hylin et al., 2013). Therefore, understanding the impact of more delicate PNN alterations on neuroplasticity is imperative for developing novel therapies targeting brain ECM. In a previous study, we developed a method allowing for the topological quantification of PNN morphology based on superresolution fluorescence microscopy (Dzyubenko et al., 2018). We showed that despite their rigidity, PNNs are subject to subtle modulation post stroke and anticipated that their transient loosening can support brain remodeling. In this work, we have further elaborated our method and investigated PNN morphology with nanoscale resolution and linked it to synaptic remodeling and neurological recovery post stroke.

## Results

### Inflammatory preconditioning reduces the infarct size but not delayed brain atrophy

We investigated post stroke brain remodeling in mice with and without stroke tolerance induced by inflammatory preconditioning. Focal cerebral ischemia was induced by transient left-sided intraluminal occlusion of the middle cerebral artery (tMCAO) for 30 minutes resulting in highly reproducible ischemic lesions located in the striatum and adjacent cortical areas, but not the motor cortex. Inflammatory preconditioning was performed by injecting 1 mg/kg LPS intraperitoneally 3 days before tMCAO (Fig. 1A), which triggered a robust peripheral immune response that we reported previously (Sardari et al., 2021). In agreement with previous studies (Marsh et al., 2009; Rosenzweig et al., 2004), the inflammatory preconditioning significantly reduced infarct volume at 7 days post ischemia (DPI), as indicated by Nissl staining quantifications (Fig 1B, C). However, preconditioning with LPS did not affect the delayed brain atrophy at 42 DPI (Fig 1B, D).

**Figure 1.**
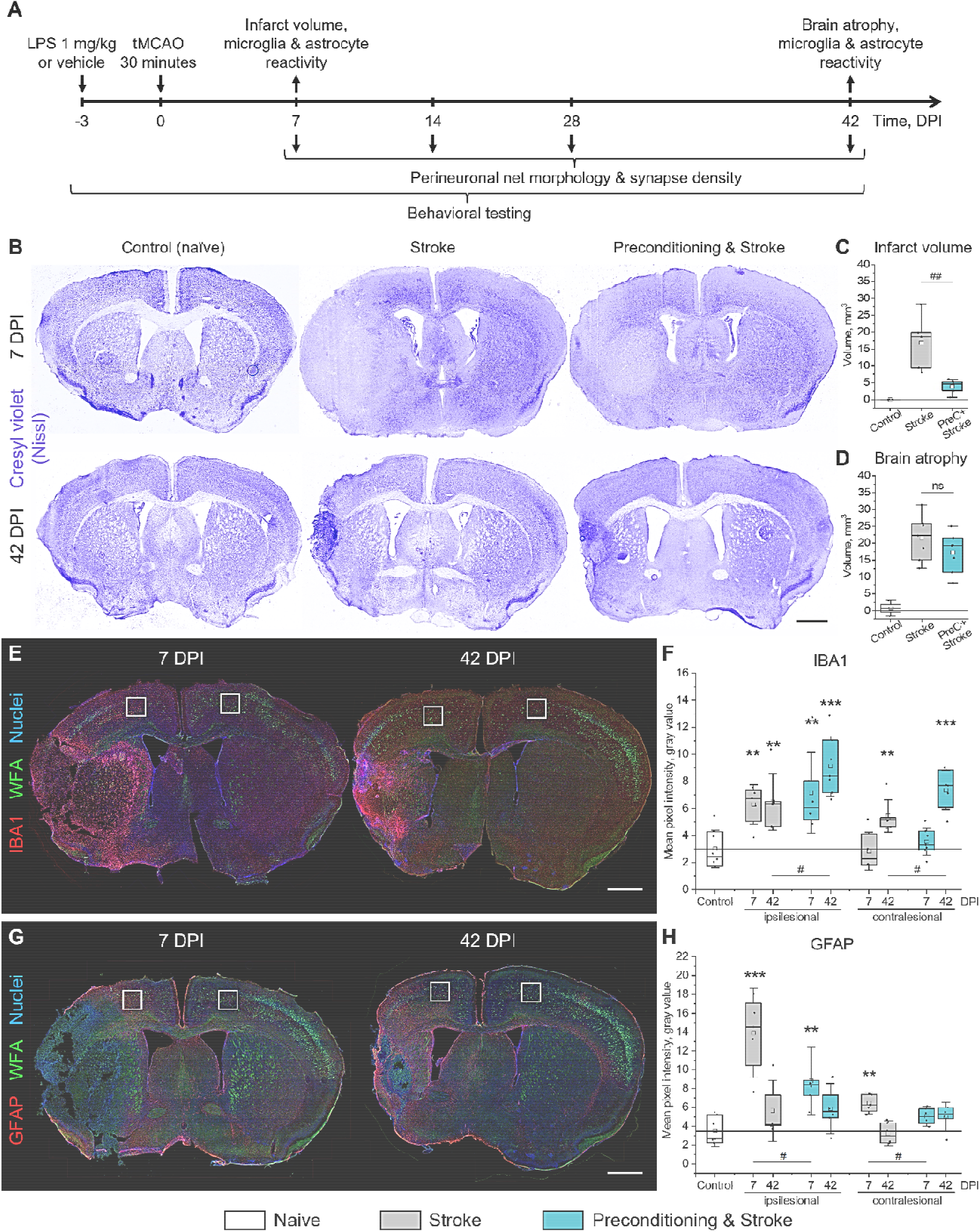
Brain damage and reactive gliosis induced by focal cerebral ischemia. (**A**) Timeline and experimental endpoints. (**B**) Cresyl violet (Nissl) staining shows focal infarcts at 7 DPI, quantified in (**C**), and brain atrophy at 42 DPI, quantified in (**D**). (**E**) Representative WFA (green) and IBA1 (red) staining at 7 and 42 DPI. (**F**) IBA1 immunoreactivity in the motor cortex L5. (**G**) Representative WFA (green) and GFAP (red) staining at 7 and 42 DPI. (**H**) GFAP immunoreactivity in the motor cortex L5. (**E, G**) White squares outline 600×600 µm regions of interest selected for analysis. Nuclei (blue) were stained with DAPI. (**C, D, F, H**) Graphs are box plots with data as dots, means as squares, medians as lines, interquartile ranges as boxes, and whiskers showing SD. Asterisks and hashes denote significant differences with the control and stroke groups, correspondingly, as indicated by two-way ANOVA and t-tests (*^,#^p < 0.05, **^,##^p < 0.01, ***p < 0.001), n = 7. Scale bars, 1 mm. DPI, days post ischemia; ns, not significant.

### Inflammatory preconditioning alters glial responses post stroke

Post-stroke reactive gliosis was evaluated using IBA1 and GFAP immunohistochemistry detecting microglia/macrophages and reactive astrocytes, correspondingly (Fig 1E, G). Focal cerebral ischemia triggered microglia/macrophage activation not only in lesion-associated areas, but also in the ipsilesional motor cortex L5, as indicated by IBA1 immunoreactivity quantification at 7 DPI (Fig. 1F). Microglia/macrophage activation persisted at 42 DPI and was also observed in the contralesional hemisphere. Inflammatory preconditioning increased IBA1 immunoreactivity at 42 DPI, but not 7 DPI. In contrast, astrocytic reactivity at 7 DPI was reduced at 7DPI in the group exposed to both preconditioning and stroke, as indicated by GFAP immunoreactivity quantification (Fig. 1H).

### Interneurons lose PNNs partially and transiently after stroke

Cortical L5 fast spiking interneurons are critical for oscillatory activity in the motor cortex, pyramidal tract output activity regulation, and motor control (Estebanez et al., 2017). These cells express parvalbumin, potassium channels Kv3.1, and are coated with a pattern of aggregated extracellular matrix forming PNNs (Hartig et al., 1999; Sigal et al., 2019). PNNs regulate several types of neuroplasticity (Carulli et al., 2020; Pizzorusso et al., 2002; Shi et al., 2019; Tewari et al., 2018), but their role in post stroke brain remodeling remains under investigated.

We analyzed the expression of PNNs around motor cortical L5 interneurons (regions of interest are outlined in Fig 1E, G) using the *Wisteria floribunda* agglutinin (WFA) that binds glycan chains of extracellular proteoglycans enriched in PNNs (Deepa et al., 2006; Haji-Ghassemi et al., 2016). Co-labelling of parvalbumin (PV) and Kv3.1 with WFA (Fig 2A, C) indicated that stroke reduced the expression of PNNs in the ipsilesional motor cortex L5 at 7 DPI (Supplementary Fig S1A). In healthy brains, 73.4±5.5% PV^+^ and 93.5±2.1% Kv3.1^+^ expressed PNNs, as evidenced by PV^+^/PNN^+^ (Fig 2B) and Kv3.1^+^/PNN^+^ (Fig 2D) cell quantifications. In mice exposed to stroke only, on average 34.3% PV^+^ and 15.3% Kv3.1^+^ neurons in the ipsilesional motor cortex lost PNN coatings during the first week post stroke. PNN expression was restored by 42 DPI. Inflammatory preconditioning reduced the transient loss of PNNs around PV^+^ neurons at 7 DP, compared with the stroke-only group (Fig 2B). In all time points and conditions evaluated, the majority of PNN^+^ cells were also PV^+^ (96%) and Kv3.1^+^ (97%), and stroke altered neither PV^+^ nor Kv3.1^+^ cell density, as shown in Supplementary Fig S1.

**Figure 2.**
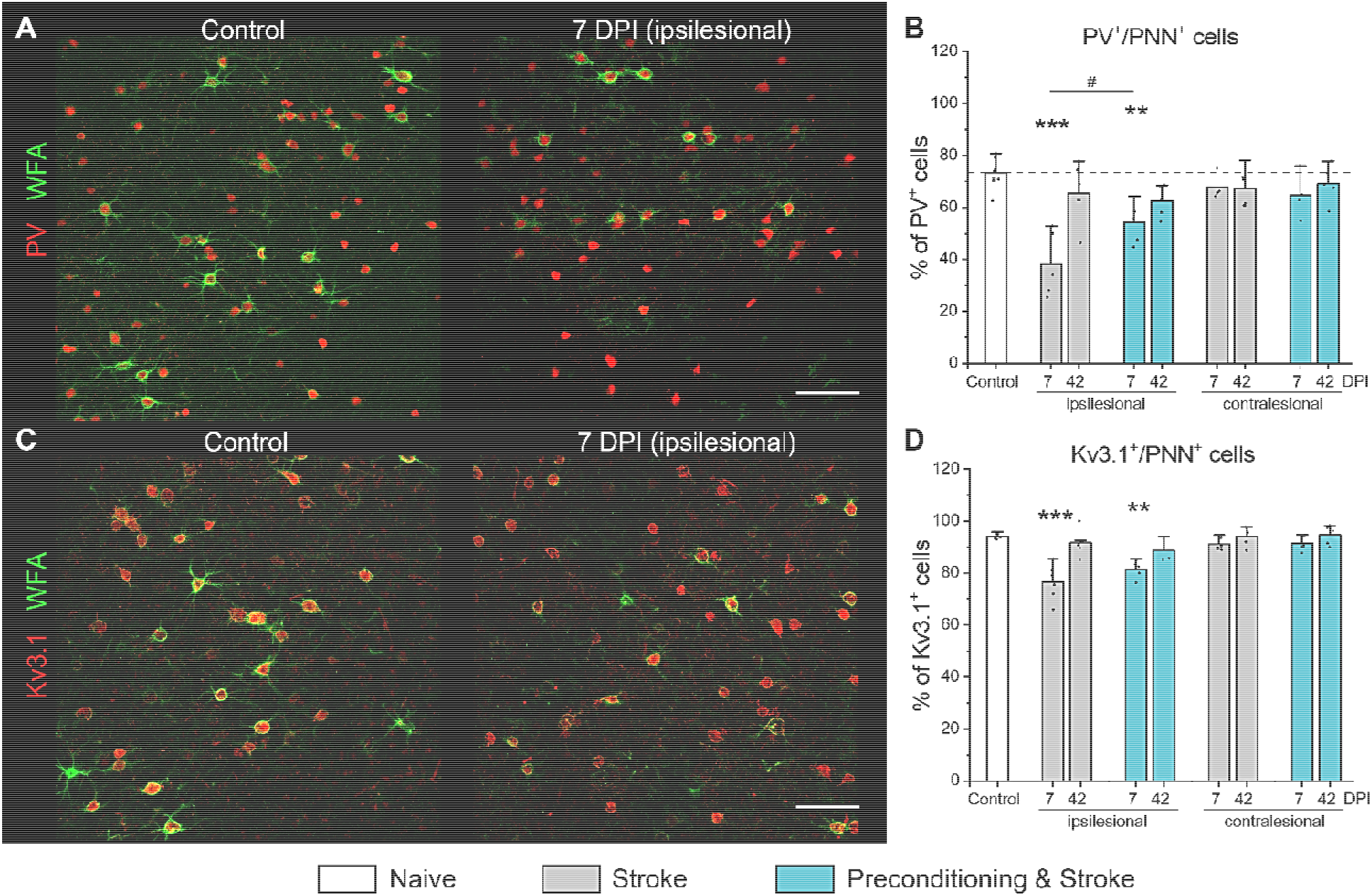
PNN expression in the motor cortex L5 post stroke. Representative immunolabeling of WFA (green) and (**A**) PV (red) or (**C**) Kv3.1 (red) shows PNN expression around motor cortical L5 interneurons. (**B, D**) Percentage of PV (**B**) and Kv3.1 (**D**) expressing neurons coated with PNNs. Graphs are bar plots showing mean±SD and data as dots. Asterisks and hashes denote significant differences with the control and stroke groups, correspondingly, as indicated by two-way ANOVA and t-tests (^#^p < 0.05, **p < 0.01, ***p < 0.001), n = 7. Scale bars, 100 µm. DPI, days post ischemia; PV, parvalbumin.

### PNN morphology analysis requires superresolution imaging

PNNs are exceptionally stable mesh-like structures in the extracellular space (ECS) consisting of densely packed proteoglycans bound together by link proteins (Carulli et al., 2010; Deepa et al., 2006). Chondroitin sulfate carrying proteoglycans of PNNs repel axons (Becker and Becker, 2002; Wang et al., 2008), and the perisomatic synapses on fast-spiking interneurons are established within the PNN facets forming synaptic pockets, which can be visualized with confocal microscopy (Fig. 3A, B).

**Figure 3.**
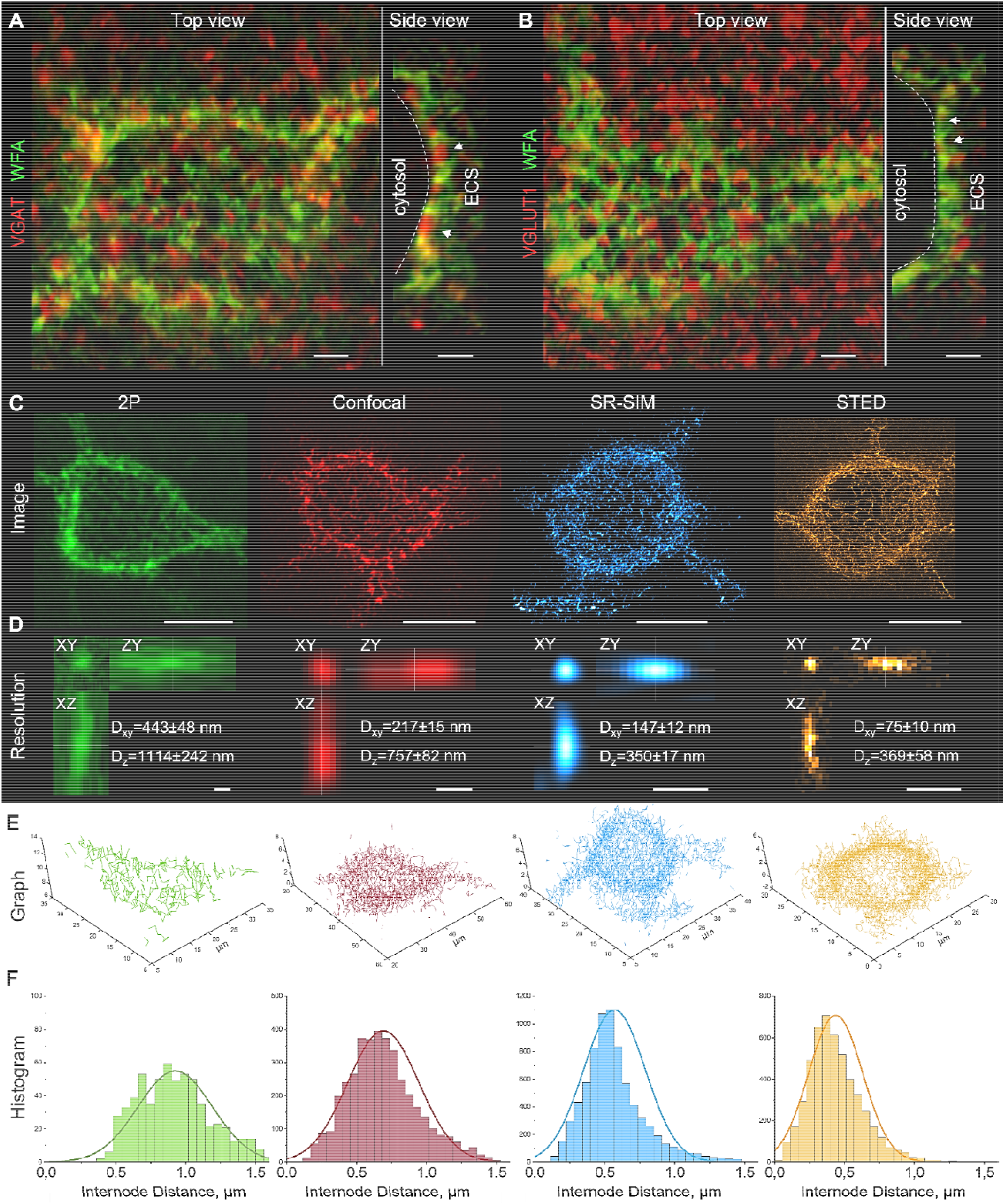
PNN morphology analysis using confocal, 2P, SIM, and STED microscopy. (**A, B**) Confocal microscopy of WFA-labeled neurons shows characteristic facet-like morphology of PNNs. PNN facets form synaptic pockets (arrowheads), within which presynaptic terminals contact the neuronal surface (dash line) and establish synapses. GABAergic (**A**) and glutamatergic (**B**) terminals are shown by vesicular GABA (VGAT, red) and glutamate (VGLUT1) transporter labeling, correspondingly. Arrowheads and dash lines indicate putative synaptic pockets and neuronal surfaces, correspondingly. Scale bars, 2 µm. (**C**) PNNs in the motor cortex L5 (control brains) were labeled with biotinylated WFA and streptavidin conjugated to Atto 490 (2P microscopy) or Star RED (confocal, SR-SIM and 3D STED microscopy) fluorophores. Images are maximum intensity z-projections. Scale bars, 10 µm. (**D**) Imaging resolution was estimated using sub-resolution microspheres (see Methods). Scale bars, 500 nm. (**E**) PNN morphology was reconstructed as graphs with nodes positioned at local fluorescence intensity maxima and edges generated by a non-redundant nearest neighbor search algorithm. (**F**) Histograms show internode distance distributions for the single PNNs shown in (**C)** and (**E**). 2P, two-photon excitation; SR-SIM, superresolution structured illumination microscopy; STED, stimulated emission depletion; D_xy_, lateral resolution; D_z_, axial resolution; ESC, extracellular space.

After stroke, PNNs undergo morphological changes beyond all-or-none degradation, and the understanding of their structural remodeling requires three-dimensional superresolution imaging (Dzyubenko et al., 2018; Sigal et al., 2019). We analyzed the morphology of PNNs using the quantitative approach that combines 3D superresolution fluorescence imaging and graph-based computational reconstruction. In brief, local fluorescence intensity maxima of PNN mesh vertices were defined as net nodes, and the internode connections (edges) were reconstructed using the non-redundant nearest neighbor search algorithm. Graphs thus obtained reflect the morphology of PNNs and allow for the quantitative analysis of their structure. In comparison to our previous report (Dzyubenko et al., 2018), we significantly improved imaging resolution by adjusting the refractive index of stained tissues (also known as tissue clearing), using fluorophores with longer emission wavelengths, and applying additional microscopy techniques.

We compared PNN morphology (Fig. 3C) visualized by four cutting-edge 3D imaging methods: two-photon (2P) excitation, confocal, superresolution structured illumination (SR-SIM) and stimulated emission depletion (STED) microscopy. Lateral (D_xy_) and axial (D_xy_) imaging resolution (Fig. 3D) was estimated as the full width at half-maximum (FWHM) using sub-resolution fluorescent beads (Ø 100 nm and Ø 40 nm for STED) embedded in the stained tissue. 2P microscopy with a small numerical aperture (NA=0.95) water immersion objective had the lowest resolution (D_xy_ = 443±48 nm, D_xy_ = 1114±242 nm) among the compared methods, resulting in the incomplete PNN graphs with long internode distances (Fig. 3E, F). Confocal, SR-SIM, and 3D STED imaging was performed using oil immersion objectives with high numerical apertures (NA=1.4 or 1.46).

The lateral resolution of confocal microscopy (D_xy_ = 217±15 nm) was very close to the diffraction limit (d = λ/2NA = 633/2.92 = 216.8 nm), but the relatively low axial resolution (D_z_ = 757±82 nm) led to the merging of closely spaced PNN nodes, resulting in imperfect graph reconstruction. Both SR-SIM and 3D STED methods had a superior resolution that was beyond the diffraction limit in all three dimensions (Fig. 3B). While SR-SIM had slightly better D_z_ (350±17 versus 369±58 nm), 3D STED significantly outperformed SR-SIM in terms of D_xy_ (75±10 versus 147±12 nm). Of note, D_z_ and D_xy_ are individually adjustable for STED and were optimized in each sample. Both SR-SIM and 3D STED imaging allowed for precise reconstruction of PNN morphology using graphs, the mathematical constructs designed for topological analysis (Fig. 3C, D).

To compare PNN morphology resolved by SR-SIM and 3D STED, we sequentially visualized the same PNN-coated neuron with the two methods (Fig. 4A, B) and aligned the obtained 3D image stacks based on the intensity maxima positions. Both methods revealed highly similar PNN structures, and both major vertices of the meshes and putative synaptic pockets were found at the same positions (Fig. 4C, D). Regions of interest showing putative synaptic pockets are magnified in Supplementary Fig. S2.

**Figure 4.**
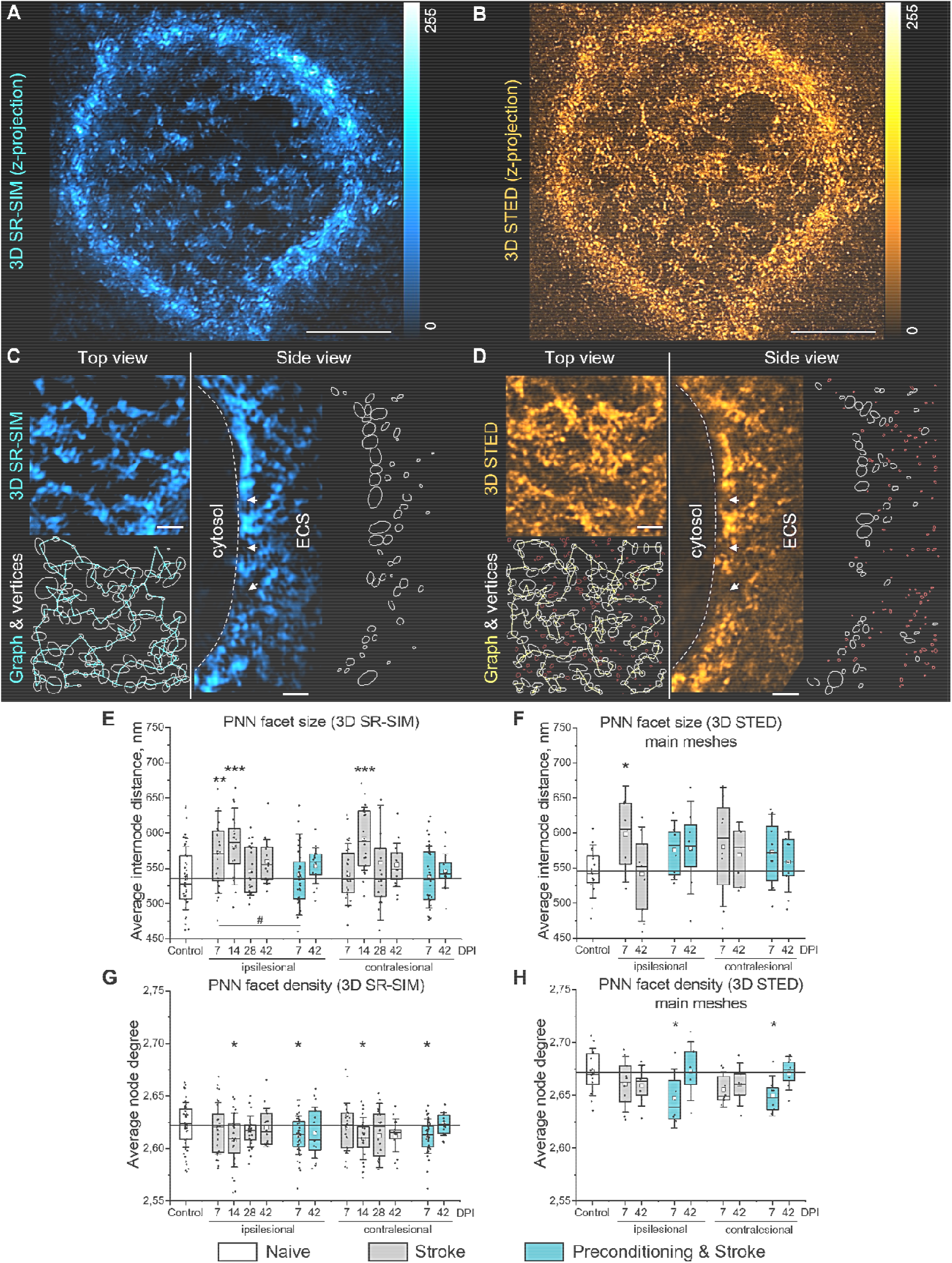
PNN morphology in the motor cortex L5 post stroke. (**A, B**) Maximum intensity z-projections show the morphology of the same PNN sequentially visualized using (**A**) SR-SIM and (**B**) STED microscopy. Scale bars, 5 µm. (**C, D**) High magnification single plane images, mesh vertices, and reconstructed graphs are shown for the same region visualized using (**C**) SR-SIM and (**D**) STED microscopy. Arrowheads and dash lines indicate putative synaptic pockets and neuronal surface, correspondingly. White outlines show PNN vertices, and red outlines are interstitial particles. Scale bars, 1 µm. (**E, F**) PNN facet size was quantified as average internode distance. (**G, H**) PNN facet density was quantified as average node degree. Graphs are box plots with data as dots, means as squares, medians as lines, interquartile ranges as boxes and SD as whiskers. Asterisks and hashes denote significant differences with the control and stroke groups, correspondingly, as indicated by two-way ANOVA and t-tests (*^,#^p < 0.05, **p < 0.01, ***p < 0.001), (**E, G**) n = 7, (**F, H**) n = 5. DPI, days post ischemia; SR-SIM, superresolution structured illumination microscopy; STED, stimulated emission depletion; ESC, extracellular space.

Due to superior lateral resolution, 3D STED microscopy detected multiple smaller vertices in addition to those detected by SR-SIM (Fig. 4D). These intensity maxima were predominantly observed at a distance more than 1 µm from the neuronal surface and seldom associated with synaptic pockets. We therefore concluded that these particles represent less condensed interstitial brain matrix and not PNNs. The small particles were filtered for PNN graph reconstruction, based on the cutoff defined by the Gaussian mixture model showing the bimodal distribution of WFA-labeled particle volumes measured by STED microscopy (Supplementary Fig. S3A, B).

### Stroke induces transient loosening of PNNs in both hemispheres

We quantified morphological changes in motor cortical L5 PNNs (regions of interest are outlined in Fig 1E, G) post stroke by measuring internode distances and node degrees (that is, the number of neighbors connected to a vertex) of the graphs reconstructing the organization of WFA-labeled meshes (Fig. 4 E-H). Average internode distance (*L)* indicates PNN facet size, and average node degree (*D*) reflects facet density.

In mice exposed to stroke only, the size of PNN facets measured with SR-SIM (Fig. 4E) was significantly increased in the ipsilesional motor cortex at 7 DPI (*L*=573±59 nm) and 14 DPI (*L*=581±55 nm), and in the contralesional hemisphere at 14 DPI (*L*=591±45 nm), compared with the control (*L*=536±45 nm). At 28 and 42 DPI, PNN facet size decreased back to control levels. Inflammatory preconditioning minimized the observed effect, and PNN facet size was not different from the control at 7 and 42 DPI in mice exposed to preconditioning and stroke. We observed only a minor decrease in the density of PNN facets (Fig. 4G) at 14 DPI in the stroke group and at 7 DPI in the preconditioning and stroke group in both ipsilesional and contralesional hemispheres. The average size of the PNN nodes was not different between the groups (Supplementary Fig. S4A).

The results obtained using SR-SIM were confirmed by 3D STED microscopy. The size of PNN facets measured with 3D STED was increased in the ipsilesional motor cortex at 7 DPI (*L*=598±60 nm versus *L*=545±33 nm in control) in mice exposed to stroke only (Fig. 4F). At 42 DPI, PNN facet size decreased back to control levels, and the size of the PNN nodes increased (Supplementary Fig. S4B), indicating condensation of ECM material. In mice with induced stroke tolerance, the facet size was not affected at 7 and 42 DPI, but we observed a minor decrease in the facet density at 7 DPI in both ipsilesional and contralesional hemispheres (Fig. 4H). In the interstitial matrix, we observed no alterations induced by stroke (Supplementary Fig. S3C-E).

### Transient PNN loosening associates with bilateral synaptic remodeling post stoke

Because PNNs have a major impact on synaptic plasticity (Fawcett et al., 2019), we further explored whether the transient loosening of PNN facets post stroke (Fig. 5A) associates with synaptic remodeling. The density of GABA- and glutamatergic terminals perforating PNNs was quantified in 33×33×5 µm regions of interest positioned in the motor cortex L5 containing a single PNN coated neuron (Fig. 5B). We quantified the local GABAergic and glutamatergic projections targeting the PNN-coated interneurons by immunolabeling the vesicular transporters of GABA (VGAT) and glutamate (VGLUT1), correspondingly.

**Figure 5.**
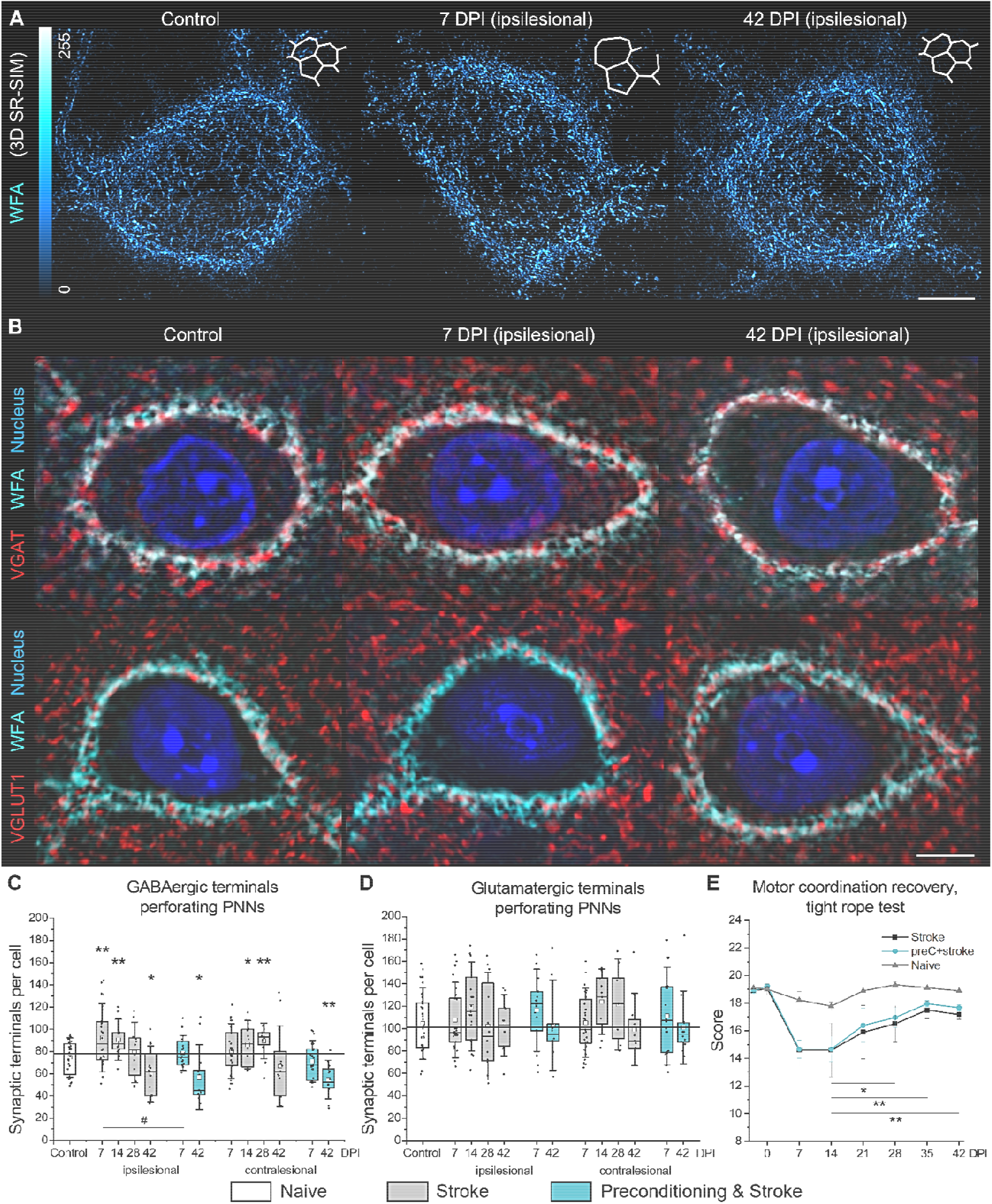
Coherent remodeling of PNNs and their perforating synaptic terminals precedes motor coordination recovery after stroke. (**A**) Representative maximum intensity z-projections show transient loosening of motor cortical PNNs after stroke detected by SR-SIM. Inlets show artistic representations of PNN facets. (**B**) Single-plane confocal images show representative immunolabeling of GABAergic axonal terminals expressing VGAT (red) and glutamatergic axonal terminals expressing VGLUT1 (red). PNNs were labeled with WFA (cyan), nuclei are shown in blue. Scale bars, 5 µm. (**C, D**) Quantifications show the number of GABAergic (**C**) and glutamatergic (**D**) terminals perforating PNNs. Graphs are box plots with data as dots, means as squares, medians as lines, interquartile ranges as boxes and whiskers showing SD. Asterisks and hashes denote significant differences with the control and stroke groups, correspondingly, as indicated by two-way ANOVA and t-tests (*^,#^p < 0.05, **p < 0.01), n = 7. (**E**) Motor coordination recovery measured by tight rope test. Data are mean±s.e.m. n = 7. DPI, days post ischemia; ns, not significant.

In healthy brains, motor cortical L5 interneurons received 75±15 (mean±SD) GABAergic inputs from their network partners, as indicated by the quantifications of VGAT terminals perforating PNNs (Fig. 5C). In mice exposed to stroke only, the number of GABAergic terminals perforating PNNs increased at 7 and 14 DPI (92±26 and 90±15 inputs per cell) in the ipsilesional motor cortex but was decreased at 42 DPI (60±21 inputs per cell). In the contralesional motor cortex, interneurons received more GABAergic inputs at 14 and 28 DPI (86±18 and 89±13 inputs per cell) but the number of VGAT terminals decreased back to control levels at 42 DPI. In mice treated with inflammatory preconditioning, the number of GABAergic terminals decreased at 42 DPI (57±24 inputs per cell) bilaterally but did not differ with the control at 7 DPI.

In healthy brains, motor cortical L5 interneurons received 104±27 (mean±SD) glutamatergic inputs from their local network partners, as indicated by the quantifications of VGLUT1 terminals perforating PNNs (Fig. 5D), and their number was not significantly altered post stoke.

### Motor recovery follows motor cortical tissue remodeling post stroke

We analyzed neurological recovery after stroke using Clark’s score for general and focal deficits and tight rope test for measuring motor performance. Neurological deficits (Supplementary Fig S5) manifested in the acute phase and decreased during the first week post stroke. Motor activity and coordination measured by the tight rope test (Fig 5E) were decreased at 7 DPI. Starting 28 DPI, we observed a gradual recovery of coordinated movements. Motor coordination was significantly restored by 42 DPI. We observed no significant influence of immune preconditioning on the motor performance recovery after stroke. These data indicate that the transient loosening of PNNs and synaptic remodeling in the motor cortex L5 precede the recovery of coordinated motor activity post stroke.

### Microglia-neuron surface contact increases post stroke

In a healthy brain, microglia cells establish direct contact with neuronal plasma membranes (Cserep et al., 2020) and can promote synaptic plasticity by remodeling extracellular matrix (Nguyen et al., 2020). In the fast-spiking PNN coated interneurons, the direct surface contact with microglia should be difficult because of the inhibitory properties of incorporated proteoglycans (Afshari et al., 2010; Dzyubenko et al., 2016; Fawcett et al., 2019). However, the transient loosening of PNNs after stroke (Fig. 5A) may facilitate microglia-interneuron interaction. We explored this possibility by quantifying the surface-to-surface contacts between microglia/macrophages labeled with IBA1 and the fast-spiking neurons expressing Kv3.1 channels (Fig. 6A) in the motor cortex L5 after stroke.

**Figure 6.**
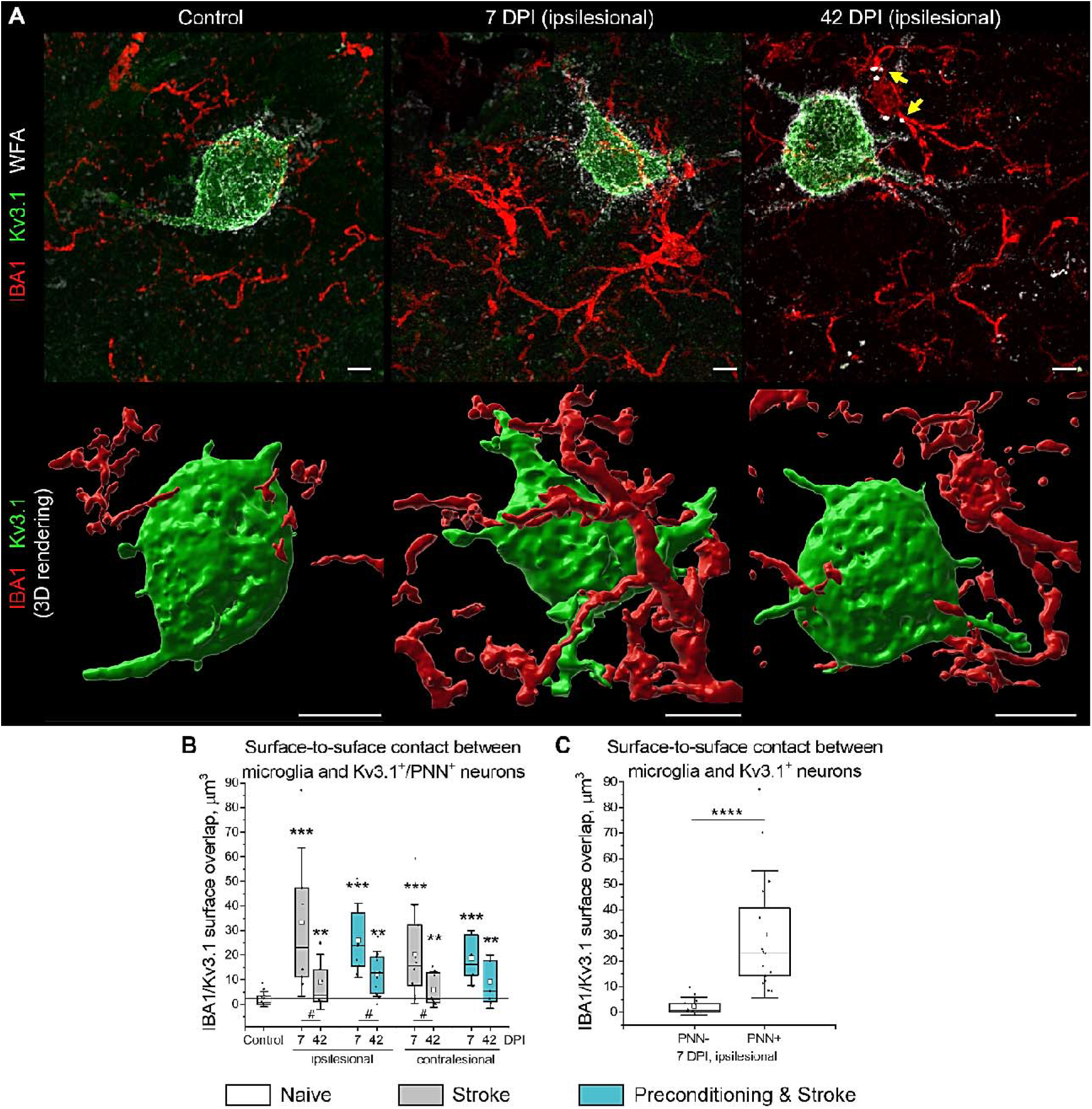
Microglia-interneuron contacts post stroke. (**A**) Confocal Z-projections show representative IBA1 (red), Kv3.1 (green) and WFA (white) immunolabeling in the motor cortex L5. Yellow arrowheads highlight speckles of WFA-labeled material inside IBA1-labelled cell bodies at 42 DPI. The corresponding 3D rendering of IBA1 and Kv3.1 surfaces is shown below. Scale bars, 5 µm. (**B**) Surface contact between microglia and Kv3.1^+^/PNN^+^ cells. (**C**) Surface contact between microglia and Kv3.1^+^ neurons with and without PNNs at 7 DPI. Graphs are box plots with data as dots, means as squares, medians as lines, interquartile ranges as boxes, and whiskers showing SD. Asterisks and hashes denote significant differences with the control and 7 DPI groups, correspondingly, as indicated by two-way ANOVA and t-tests (^#^p < 0.05, ***p < 0.001, ****p < 0.00001). n = 5. DPI, days post ischemia.

Under all experimental conditions that we investigated, multiple microglia processes were observed in close proximity to neuronal membranes expressing Kv3.1, and at least two microglia cell bodies were present within a 50 µm radius around the soma of every Kv3.1^+^ neuron. While in the healthy brains, the direct contacts between microglia and Kv3.1^+^/PNN^+^ neurons were point-like and not numerous, the IBA1/Kv3.1 surface overlap strongly increased at 7 DPI in both ipsilesional and contralesional hemispheres (Fig. 6B). At 42 DPI, microglia-interneuron contact surface decreased in comparison to 7 DPI but remained significantly larger than in control. In mice with inflammatory preconditioning, the IBA1/Kv3.1 surface overlap was increased similar to the stroke only group.

As demonstrated by 3D surface rendering (Fig. 6A), microglia enwrapped significant parts of neuronal surfaces at 7 DPI. Interestingly, the extensive contacts between Kv3.1^+^ neuronal membranes and microglial processes were observed only on Kv3.1^+^/PNN^+^ cells, indicating the high preference of microglia to contacting PNN-coated neurons (Fig. 6C). In addition, we observed multiple WFA-labeled speckles inside IBA1-labeled cells at 42 DPI (Fig. 6A), suggesting that microglia/macrophages can phagocyte PNN components after stroke.

## Discussion

We herein demonstrate that the increased size and reduced density of PNN facets associate with synaptic reorganization preceding the recovery of motor coordination after stroke. Noteworthy, morphological changes in motor PNNs revealed by SR-SIM were confirmed using 3D STED microscopy. The coherent remodeling of PNN structure and GABAergic axonal terminals on motor cortical L5 interneurons suggests a novel mechanism of stroke recovery that involves ECM modulation in both ipsi- and contralesional hemispheres. During the subacute stroke phase at 7 DPI, PNN loosening correlates with the increasing number of perforating axonal terminals expressing VGAT, which agrees with increased GABAergic phasic activity during the first week post stroke (Hiu et al., 2016). In the chronic stroke phase at 42 DPI, PNN morphology returns back to norm, but the number of GABAergic inputs received by motor cortical L5 interneurons is significantly reduced. We hypothesize that these dynamic changes in motor cortical inhibitory connectivity arise from the tripartite interaction between PNNs, synapses, and microglia.

PNNs cover the soma, proximal dendrites and initial axonal segments of the fast-spiking interneurons that express calcium binding protein PV and potassium channels Kv3.1 with fast activation and deactivation kinetics (Hartig et al., 1999; Matsuda et al., 2021). By creating facet-like structures, PNNs compartmentalize neuronal surface and restrict synapse formation to the areas devoid of inhibitory CSPGs that repel axons (Becker and Becker, 2002). The three-dimensional organization of PNN facets resembles wells that are approximately 1 µm deep, with neuronal plasma membrane at the bottom and opening towards the extracellular space. Because most of these wells are occupied by perforating axonal terminals, we here propose to call them synaptic pockets. After stroke, the increased size of synaptic pockets allows for new synapse formation. Recent findings show that synapses continuously wane and re-emerge *in vivo* and that GABAergic terminals are especially dynamic with about 60% of them retracting and returning within a few days (Villa et al., 2016). We propose that the loosening of PNNs creates larger permissive regions supporting GABAergic synapse plasticity and results in increased GABAergic input to fast-spiking interneurons at 7 and 14 DPI. These new synapses are not stabilized in the long term though, which leads to the decreased number of VGAT terminals perforating PNNs at 42 DPI. Importantly, we observed PNN loosening and synaptic remodeling in both ipsi- and contralesional hemispheres, which suggests that ECM modulation supports bilateral rewiring of brain circuits after stroke.

The removal of new GABAergic synapses that are established during the post-acute stroke phase is likely mediated by activated microglia. Recent evidence indicates that microglia cells can selectively sculpt inhibitory connectivity (Favuzzi et al., 2021) by eliminating presynaptic terminals using a phagocytic mechanism known as trogocytosis (Weinhard et al., 2018). Here, we observed that every PNN-coated fast-spiking interneuron in the motor cortex L5 is always adjacent to a few highly ramified microglia cells. Our data shows that although increased IBA1 immunoreactivity persists in the chronic stroke phase, the surface-to-surface contact between interneurons and microglia is more extensive at 7 DPI than at 42 DPI. Interestingly, the surface of microglia-interneuron contacts on PNN^+^ cells is 30 times larger than on PNN^-^ neurons at 7DPI, indicating the high microglial preference for enwrapping PNN-coated neurons in the ischemic brain.

At 42 DPI, we observed speckles of WFA-labeled substance within IBA1-labeled cell bodies, suggesting that microglia activation contributes to PNN loosening at 7 DPI. Notably, activated microglia facilitate the loss of PNNs in rodent models of Alzheimer’s (Crapser et al., 2020b) and Huntington’s (Crapser et al., 2020a) diseases. We suppose that after stroke, microglia attenuate plasticity inhibiting PNN properties in the early post-acute phase, allowing for the formation of additional GABAergic synapses on motor cortical L5 interneurons. Noteworthy, inhibitory interneurons expressing parvalbumin and PNNs play a major role in regulating activity and synchronization in neuronal networks (Favuzzi et al., 2017). The increased inhibitory input to inhibitory interneurons can switch the excitation-inhibition balance in motor cortical microcircuits towards excitation, thereby promoting motor coordination recovery that we observed starting 28 DPI. During the chronic stroke phase, PNNs recover completely and restrict new synapse formation, while the non-stabilized inputs are removed by microglia. Thus, PNNs and microglia negatively regulate inhibitory input to inhibitory interneurons, which can limit functional recovery in the chronic stroke phase.

Admittedly, our hypothesis is based on correlations and needs further validation using intravital superresolution imaging. Here, we revealed the transient remodeling of PNNs in the motor cortical L5 after stroke using SR-SIM and confirmed this effect using STED microscopy. While SR-SIM can be applied for multi-label imaging in relatively large volumes, it requires sophisticated computational processing that may generate artifacts (Heintzmann and Huser, 2017). STED microscopy has superior resolution compared to SR-SIM and does not generate any processing-related artifacts (Berning et al., 2012). However, STED microscopy is more challenging to perform in multi-color modes and commonly uses short-distance objectives that make 3D imaging in large volumes limited.

In this work, we visualized and quantified synaptic terminals and microglial-neuron contacts using high-resolution confocal microscopy. Nevertheless, our measurements relied on immunohistochemical procedures incompatible with intravital imaging. In the future, the tripartite interaction between PNN, presynaptic terminals, and microglia post stroke can be verified using reversibly switchable fluorescent proteins and multi-label *in vivo* STED microscopy (Willig et al., 2021). In comparison to our previous work (Dzyubenko et al., 2018), here we improved the method for PNN morphology reconstruction by significantly improving imaging resolution and optimizing the graph computation algorithm. As a result, the internode distances that we measured in this study are twice smaller than detected with our previous method. Improved resolution of imaging allowed us to detect the previously unnoticed increase of PNN facet size after stroke. With the earlier method, we detected a slight decrease in average internode distance, which is misleading and due to the merging of the adjacent PNN nodes because of insufficient resolution. Similar to our previous study, here we detected a decrease in the average degree of PNN nodes after stroke. However, the correct connectivity degrees for PNN nodes are between 2.5 and 2.8 as we show here, and not between 2.5 and 4 as we previously reported due to insufficient imaging resolution. Conclusively, superior microscopy techniques and cross-validation of imaging approaches in this study allowed for the detection of correct parameters of PNN morphology. Using the improved methodology, we revealed that the structural alterations in PNNs are associated with synaptic remodeling and functional recovery post stroke.

The superior resolution of STED microscopy and exceptional photostability of WFA labeling allowed for the detection of interstitial ECM particles that did not associate with PNNs. In rodent models of learning and memory formation, the interstitial matrix regulates axonal sprouting and synaptic input density, while PNNs control the number of synaptic spines and receptor mobility (Fawcett et al., 2022). In the stroke model we used herein, the interstitial matrix density was not affected, and the dynamic modulation of GABAergic input density on motor cortical L5 interneurons was associated with structural rearrangements in PNNs. This evidence indicates that the role of cortical PNNs in post stroke recovery differs from the memory-related function of hippocampal PNNs.

Despite the prominent changes in PNN morphology, the number of glutamatergic inputs on the fast-spiking motor cortical L5 interneurons is not affected post stroke. Noteworthy, the glutamatergic terminals that we detected here by VGLUT1 immunolabeling represent the local excitatory input within motor cortical microcircuits and not the thalamocortical afferents (Fremeau et al., 2004). The higher stability of glutamatergic inputs on interneurons post stroke may involve additional mechanisms independent of PNNs (Benson and Huntley, 2012; Van Horn and Ruthazer, 2019), which agrees with our recent study showing that the depletion of ECM primarily affects inhibitory and not excitatory synapses (Dzyubenko et al., 2021).

Here, we compared PNN morphology alterations, synaptic remodeling, and reactive gliosis after focal cerebral ischemia in mice with and without induced stroke tolerance. Inflammatory preconditioning via systemic injection of LPS has been shown to reduce the severity of stroke in animal models (Marsh et al., 2009; Stevens et al., 2014). Our data indicate that exposure to inflammatory stress before stroke reduces infarct volume, attenuates PNN loosening, and prevents synaptic alterations during the subacute stroke phase. Therefore, the extent of structural changes in PNNs and their perforating GABAergic synapses depends on the severity of ischemic injury. In the chronic phase though, GABAergic input is similarly reduced in mice with and without induced stroke tolerance. Inflammatory preconditioning does not reduce brain atrophy and even increases microglial reactivity at 42 DPI. We also observed no significant effect of inflammatory preconditioning on motor recovery in the chronic stroke phase, which calls into question the translational value of this approach for improving long-term recovery after stroke.

Our results suggest that the alternating morphology and integrity of PNNs in the motor cortex can be harnessed for promoting neurological recovery in the chronic stroke phase. While the intrinsic brain remodeling post stroke involves only transient loosening of PNNs, prolonging the partial PNN degradation during the post-acute period can extend the opening neuroplasticity window into the chronic stroke phase. In addition, modulation of CSPG sulfation is a promising target for improving post stroke rehabilitation. The plasticity-inhibiting properties of CSPGs depend on the ratio between 4-sulfated and 6-sulfated disaccharides (Galtrey and Fawcett, 2007; Wang et al., 2008). Outgrowing axons avoid 4-sulfated CSPGs, and their selective cleavage with arylsulfatase B has been proposed for promoting nerve regeneration (Pearson et al., 2018). Conclusively, exploring the possibilities for modulating the chemical composition and structure of cortical PNNs is a novel and promising target for post-stroke neuroplasticity research.

## Acknowledgments

The authors sincerely appreciate the grant support by the German Research Foundation (DFG, projects 389030878 and 405358801, to DMH, project 467228103 to ED). For the maintenance of equipment and advice on microscopy applications, the authors are especially thankful to the IMCES (imaging center Essen) staff.

## Author Contributions

E.D. and D.M.H. designed and planned the study. D.Y. and M.S. performed animal surgeries and coded the experimental groups to blind other experimenters. D.Y. conducted animal behavior tests. E.D., E.T, P.L and B.S. performed immunocytochemistry and SR-SIM imaging. E.D. and K.W. performed STED imaging and analyzed data. E.D. performed widefield, confocal, 2P microscopy, and associated analyses. E.D., K.W., and D.M.H. drafted the manuscript. All authors discussed the data and contributed to the final version of the manuscript.

## Methods

### Legal issues, animal housing, and randomization

Experimental procedures were conducted in accordance with European Union (Directive 2010/63/EU) guidelines for the care and use of laboratory animals and approved by the local government (Bezirksregierung Düsseldorf). C57BL/6j mice were kept in groups of 5 animals per cage, inverse 12/12 h light/dark cycle, and access to food and water *ad libitum*. All efforts were made to reduce the number of animals in the experiments. The groups were randomly assigned using dummy names, and the experimenters were blinded to group coding during sample preparation, data acquisition, and analysis.

### Cerebral ischemia, stroke tolerance induction, and tissue collection

Wildtype male C57/Bl6 mice at the age of 2 months were randomly assigned into three groups: stroke, preconditioning and stroke, and naive control. Each group included 7 animals. Focal cerebral ischemia was induced by transient left-sided intraluminal middle cerebral artery occlusion (tMCAO) for 30 min as described previously (Dzyubenko et al., 2022). In brief, mice were anesthetized with 1.5% v/v isoflurane (carrier gas was N_2_O with 30% v/v O_2_) and 150 µl of buprenorphine was injected subcutaneously. After exposing and ligating the lower part of the left common carotid artery (CCA), a silicon-coated nylon monofilament was introduced through a fine incision and advanced until the bifurcation of the middle cerebral artery (MCA). Cessation of blood supply in the MCA territory was verified by measuring laser Doppler flow (LDF). After 30 minutes, the filament was removed to induce reperfusion, which was controlled by LDF recording. The tMCAO procedure resulted in highly reproducible ischemic lesions located in striatum and adjacent cortical areas, but not motor cortex and the produced infarcts had similar volumes as in our previous studies (Dzyubenko et al., 2018; Sardari et al., 2021).

Stroke tolerance was induced by inflammatory preconditioning with 1 mg/kg LPS that was injected intraperitoneally 3 days before tMCAO to trigger robust peripheral immune response that we reported previously (Sardari et al., 2021). Experimental endpoints were defined as 7-, 14-, 28-, and 42-days post ischemia (DPI) to evaluate post stroke brain remodeling during post-acute and chronic stroke phases (Fig. 1A). Upon reaching the pre-defined endpoints, animals were anesthetized with 100 µl of ketamine-xylazine (1:3) and sacrificed by transcardiac perfusion with 4% paraformaldehyde (PFA) in normal saline. The brains were removed and immersed in 4% PFA solution for 12 hours at 4°C. Tissues were cryoprotected in sucrose gradient solutions (10-30%), carefully dried, frozen, and stored at -80 °C until further processing.

### Infarct volume and brain atrophy measurement

Coronal sections of the brain (20 µm thick) were collected at 500 µm intervals across the forebrain using a Leica CM1950 cryostat and placed onto cold microscope slides (ThermoFisher Scientific, Cat# J1800AMNT). The sections were stained with cresyl violet (that is, Nissl) and scanned using the AxioObserver Z1 microscope (objective Plan-Apochromat 10×/0.45 M25; Zeiss, Jena, Germany). The infarct volume (IV) was measured at 7 DPI as IV = Σ(IA*Δ), where IA is the infarcted area on a section and Δ is the interval between sections. Brain atrophy at 42 DPI was determined by subtracting the areas of surviving tissue in the ipsilesional hemisphere from the area of the contralesional hemisphere. Atrophy volume (AV) was calculated as AV = Σ(AA*Δ), where AA is the atrophy area on a section and Δ is the interval between sections.

### Neurological deficits and motor performance tests

General and focal neurological deficits were analyzed using Clark’s neuroscore (Clark et al., 1997) daily until 7 DPI, every 3 days until 14 DPI, and weekly until 42 DPI. Post stroke recovery of motor activity and coordination was assessed by tight rope at baseline, 7, 14, 21, 28, 35, and 42 DPI as described previously (Sardari et al., 2021). In brief, the tight rope test measures the time until the animals reach the platform from the middle of a 60-cm-long rope. Mice were pre-trained over 1–2 days before MCAO ensuring that they were able to reach the platform within ten seconds.

### Immunohistochemical procedures

Free-floating coronal sections (30 μm thick) were obtained at the level of bregma +0.5 to +1 mm using a Leica CM1950 cryostat and stored until use at -20 °C in 1:1 mixture of phosphate buffer saline (PBS) and ethylene glycol with 1% polyvinyl pyrrolidone. For immunohistochemistry, the sections were rinsed 0.1 M PBS and permeabilized with 0.3% w/v Triton X-100 in PBS. Non-specific antibody binding was blocked by applying a mixture of 10% v/v ChemiBLOCKER (Cat# 2170, Millipore, Burlington, MA, U.S.A.) and 5% v/v normal donkey serum in PBS for 12 hours at room temperature with gentle agitation. Sections were incubated with primary antibodies for 48 hours at 4 °C in PBS with 0.01% w/v Triton X-100. Astrocytes and microglia/macrophage cells were labeled using rat anti-GFAP (1:300; Cat# 13-0300, Thermo Fisher Scientific, Waltham, MA, U.S.A) and rabbit anti-IBA1 (1:500; Cat# 019-19741, Wako, Neuss, Germany) or guinea pig anti-IBA1 (1:300, Cat# 234308, Synaptic Systems, Göttingen, Germany) antibodies. Interneurons were detected with rabbit anti-Parvalbumin (1:500; Cat# 195002, Synaptic Systems) and rabbit anti-Kv3.1b (1:1000, Cat# APC-014, Alomone Labs, Jerusalem, Israel). PNNs were labeled with biotinylated WFA (1:100, B-1355, Vector Laboratories, Burlingame, USA). Glutamatergic and GABAergic synaptic terminals were detected using guinea pig anti-VGLUT1 (1:500, Cat# 135304, Synaptic Systems) and guinea pig anti-VGAT (1:500, Cat# 131103, Synaptic Systems) antibodies. For fluorescence detection, we used secondary antibodies conjugated to Alexa 488, 594 and 647 or streptavidin conjugated to Atto 495, Atto 590 or Abberior STAR RED dyes. Nuclei were counterstained with DAPI (1:1000, D1306, ThermoFisher). The bound antibodies were stabilized by incubating the sections in 2% w/v PFA for 30 min at room temperature. For high-resolution imaging with confocal, SR-SIM and STED microscopy with oil immersion objectives, the refractive index of stained tissues was adjusted to 1.5 using 2,2’-thiodiethanol (TDE, Cat# 166782, Merck, Darmstadt, Germany), which is widely used for tissue clearing (Costantini et al., 2015).

### Quantification of glial and neuronal markers

The expression of GFAP and IBA1 markers was analyzed using a AxioObserver Z1 microscope (objective Plan-Apochromat 10×/0.45 M25; Zeiss). In the whole-section images obtained by tiling, 600×600 µm regions of interest (ROIs) were selected in the motor cortical layer 5 as shown in Fig 1E, G, and the mean pixel intensity was measured using ImageJ (National Institutes of Health, Bethesda, MD, U.S.A.). In each animal, four images obtained from two adjacent brain sections were analyzed in the ipsilesional and contralesional motor cortex.

Cell density of interneurons expressing PV, Kv3.1 and PNNs was quantified manually in 600×600×10 µm ROIs obtained in the motor cortical L5 regions using the LSM 710 confocal microscope (Zeiss, 20x Plan Apochromat objective, NA 0.8, pixel size 0.21 µm). In each animal, four image stacks obtained from two adjacent brain sections were analyzed in the ipsilesional and contralesional motor cortex.

### Quantification of synaptic inputs and microglia-neuron surface contact

The number of axonal terminals perforating PNNs was quantified in 33×33×5 µm z-stacks obtained with high-resolution confocal microscopy using the LSM 710 microscope (Zeiss, 100x alpha Plan-Apochromat objective, NA 1.46, voxel size 60×60×500 nm). ROIs were positioned in the motor cortex L5 containing a single PNN coated neuron. Synaptic terminals expressing VGAT or VGLUT1 that associated with WFA labeling were counted using an automated ImageJ routine (see Supplementary Code 1).

The surface of microglia-neuron contacts was quantified in 75×75×10 µm z-stacks obtained with high-resolution confocal microscopy using the LSM 710 microscope (Zeiss, 63x alpha Plan-Apochromat objective, NA 1.4, voxel size 70×70×450 nm). ROIs were positioned in the motor cortex L5 containing a single PNN coated neuron. Surfaces representing IBA1 (microglia) and Kv3.1 (fast-spiking interneurons) labeled cells were generated by automated thresholding with IMARIS 9.9 software (Oxford Instruments, Stockholm, Sweden) using the standard Surfaces function. Area of contact between cells was quantified as the intersection between IBA1 and Kv3.1 surfaces.

### Microscopy techniques and resolution measurements

The morphology of WFA-labeled PNNs was analyzed using the four 3D imaging methods: two-photon excitation (2P), confocal, superresolution structured illumination (SR-SIM), and 3D stimulated emission depletion (STED) microscopy.

2P microscopy of PNNs labeled with Atto 495 was performed using a Leica TCS SP8 (25x HCX IRAPO L 25x water immersion objective, NA 0.95) microscope. ROIs (92×92×11.5 µm, voxel size 90×90×480 nm) were scanned as z-stacks. Excitation laser (Titanium:Sapphire Chameleon Vision II) was tuned to 940 nm, and the output power was 1.712 W. Emitted fluorescence (500 – 570 nm detection wavelength) was collected using a hybrid detector. Imaging resolution estimated by measuring full width at half maximum (FWHM) of Ø 100 nm TetraSpeck microspheres (Cat# T7279, Thermo Fisher Scientific) using the same microscope settings as for PNN imaging.

Confocal microscopy of PNNs labeled with Atto 590 was performed using a LSM 710 (100x alpha Plan-Apochromat objective, NA 1.46) microscope. ROIs (65.4×65.4×5.5 µm, voxel size 60×60×350 nm) were scanned as z-stacks. Excitation laser (561 nm DPSS) was used at 10% of maximum output power to reduce photobleaching. Confocal pinhole size (585 – 655 nm detection wavelength) was adjusted to 1 Airy unit. Imaging resolution was estimated by measuring FWHM of Ø 100 nm TetraSpeck microspheres (Cat# T7279, Thermo Fisher Scientific) using the same microscope settings as for PNN imaging.

SR-SIM microscopy of PNNs labeled with Atto 590 or Abberior Star RED was performed using a Carl Zeiss Elyra PS.1 (100x alpha Plan-Apochromat objective, NA 1.46) microscope. ROIs (49.4×49.4×6.7 μm, voxel size 20×20×100 nm) were scanned as z-stacks. Excitation lasers (561 and 642 nm OPSL) were used at 5% of maximum output power to reduce photobleaching. We used the 51 µm grid (5 rotations, 5 phases) for 585 – 655 nm and 649 – 755 nm detection wavelengths. The output superresolution images were computed using the automatic processing mode using the Zen Black software (Zeiss). Imaging resolution was estimated by measuring FWHM of Ø 100 nm TetraSpeck microspheres (Cat# T7279, Thermo Fisher Scientific) using the same microscope settings as for PNN imaging.

STED microscopy of PNNs labeled with Abberior Star RED was performed using the custom-built setup (Wegner et al., 2017) at the Max Planck Institute for Multidisciplinary Sciences in Göttingen, Germany. We used an oil immersion objective (HCX PL APO 100x/1.40 OIL STED, Leica Microsystems). ROIs (25×25×5 μm, voxel size 30×30×100 nm) were scanned as z-stacks. Pixel dwell time was 10 µs. Pulsed excitation light of 650 nm was spectrally filtered from a white light source (Wegner et al., 2017) and applied with an average power of 2.7 µW at the back focal plane of the objective. The STED laser (Katana 08 HP, OneFive GmbH, Regensdorf, Swiss), providing nanosecond pulses at 775 nm, was employed with a power of 221 mW. The STED beam was shaped by a spatial light modulator (Abberior Instruments) with a 2π-vortex and π phase delay for x/y- and z-depletion, respectively. The x/y-lateral and z-axial resolution was independently adjusted and estimated by measuring the FWHM of Ø 40 nm TransFluoSpheres (Ex/Em 633/720 nm, Cat# T8870, Thermo Fisher Scientific) using the same microscope settings as for PNN imaging.

### Reconstruction and analysis of PNN morphology

Structural organization of PNNs was visualized by superresolution microscopy and their morphology was analyzed using the quantitative graph-based computational reconstruction. The previously reported method (Dzyubenko et al., 2018) was optimized to improve imaging resolution and reliability of quantifications. The new optimized MATLAB code is provided here as Supplementary Code 2. In each animal, a total of 8 individual PNNs were analyzed in the ipsilesional and contralesional motor cortex. In brief, the 3D image stacks obtained by superresolution STED or SR-SIM microscopy were imported into the IMARIS 9.9 program (Oxford Instruments) and local fluorescence intensity maxima were defined as net nodes representing PNN mesh vertices. The internode connections (edges) were reconstructed using the non-redundant nearest neighbor search algorithm in MATLAB. The resulting graphs were used to derive morphological metrics that characterize the structure of PNN facets. More specifically, we quantified average internode distances (*L)* indicating PNN facet size and the average node degrees (*D*) reflecting facet density.

### Statistical analysis

All quantitative data was presented as box plots indicating the mean (empty square), the median (line), 25-75 % range (borders) and SD (whiskers) of data distribution. For all datasets, the normality of distribution was analyzed using the Kolmogorov-Smirnov test. The differences between groups were evaluated using one-way (infarct volume and brain atrophy) or two-way (all other readouts) analysis of variance (ANOVA) and post-hoc pairwise t-tests. For multiple comparisons, Bonferroni corrections were applied.

## Supplemental information

**Figure S1.**
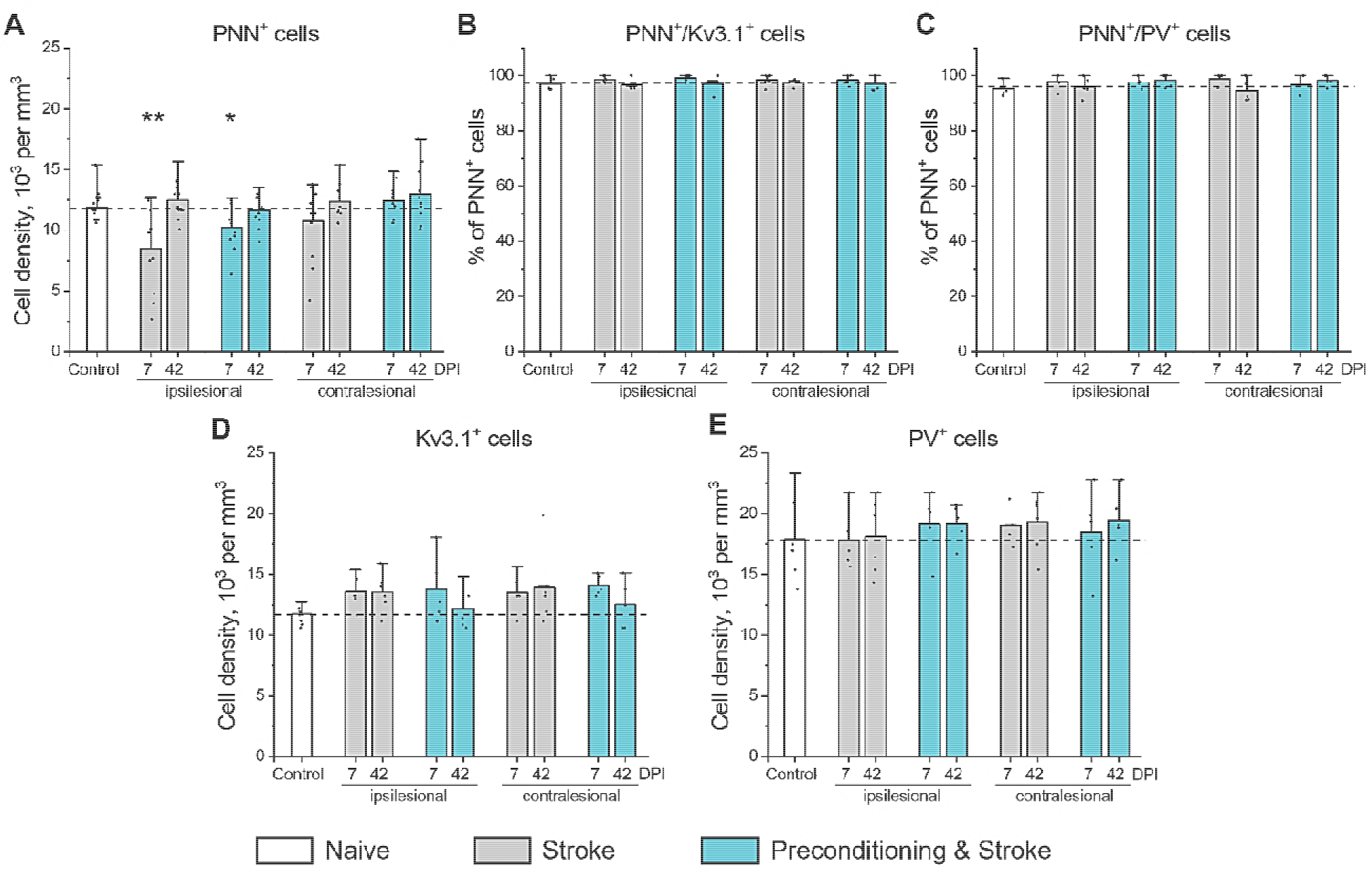
Expression of Kv3.1 and parvalbumin in the motor cortical PNN^+^ neurons. (**A**) Cell density of neurons expressing PNNs. (**B**) Percentage of PNN^+^ neurons expressing Kv3.1. (**C**) Percentage of PNN^+^ neurons expressing PV. (**D, E**) Cell density of neurons expressing PV (**D**) and Kv3.1 (**E**). Graphs are bar plots showing mean±SD and data as dots. Asterisks and hashes denote significant differences with the control group, as indicated by two-way ANOVA and t-tests (*p < 0.05, **p < 0.01), n = 7. DPI, days post ischemia; PV, parvalbumin.

**Figure S2.**
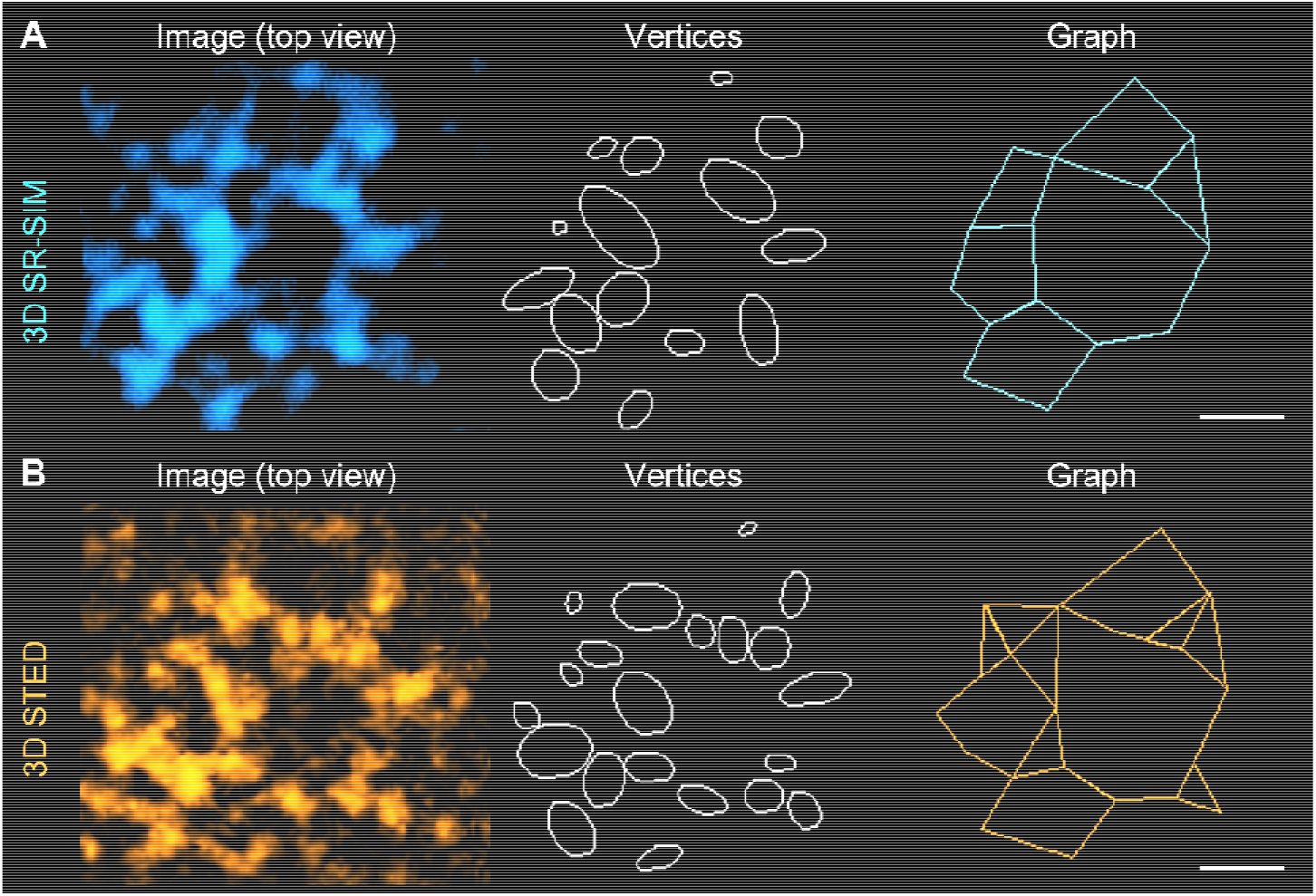
Structure of a putative synaptic pocket. High magnification single plane images, mesh vertices, and the reconstructed graphs are shown for the same region visualized using (**A**) SR-SIM and (**B**) STED microscopy. Scale bars, 500 nm.

**Figure S3.**
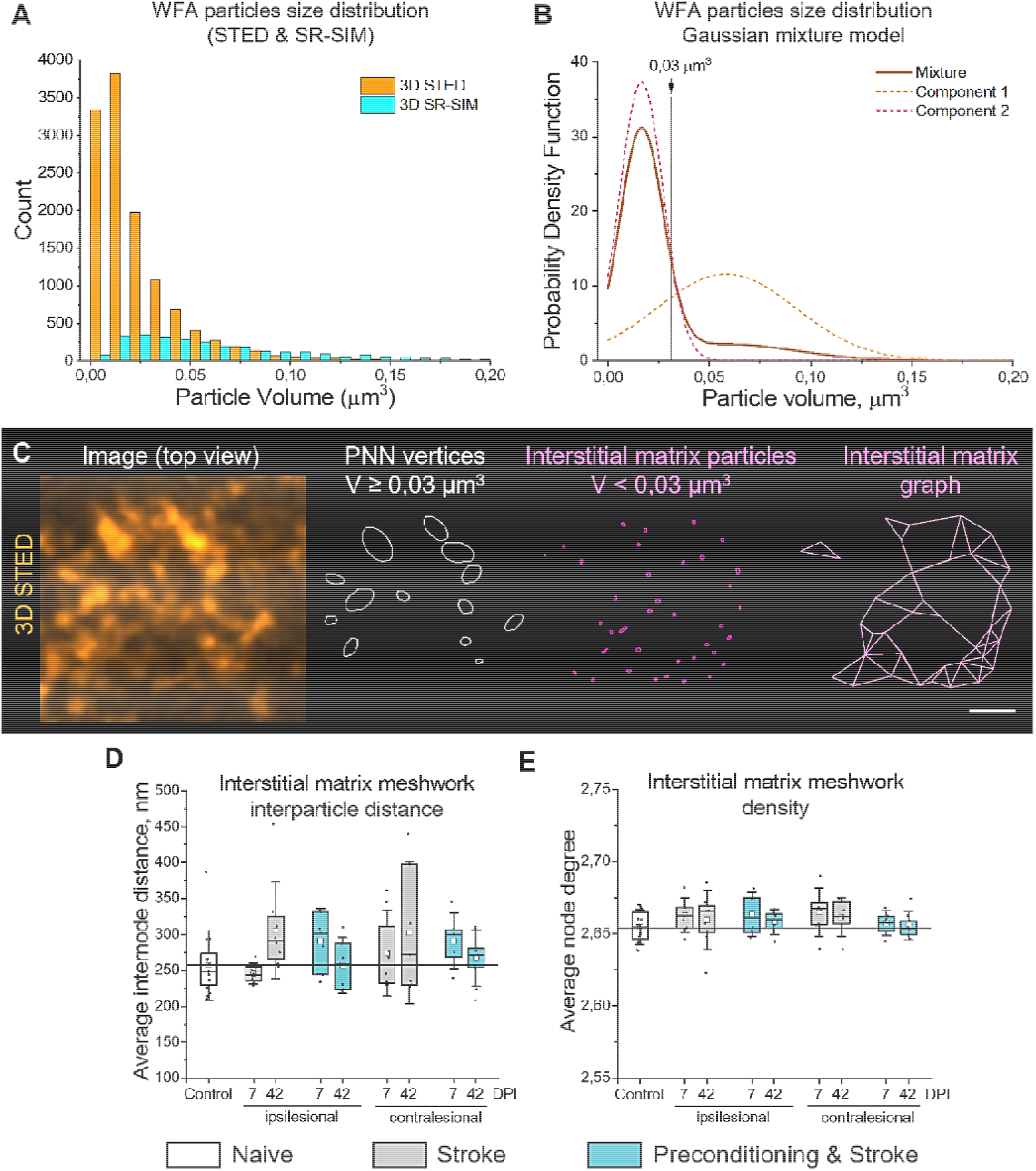
STED microscopy detects both PNN vertices and interstitial matrix particles. (**A**) Histograms show the distribution of WFA-labeled particle volumes measured with SR-SIM and STED microscopy in control mice (n=5). Bin size, 0.01 µm^3^. (**B**) Gaussian mixture model indicates the bimodal distribution of PNN vertex volumes measured by STED microscopy. The 0.03 µm^3^ cutoff was chosen to filter the small vertices not associating with PNNs. (**C**) High magnification single plane STED image, PNN vertices, interstitial matrix particles, and the reconstructed interstitial matrix graph are shown. Scale bars, 500 nm. Interparticle distance (**D**) and meshwork density (**E**) quantifications indicate no significant alterations in the interstitial matrix post stroke. Graphs are box plots with data as dots, means as squares, medians as lines, interquartile ranges as boxes and whiskers showing SD. n=5. DPI, days post ischemia.

**Figure S4.**
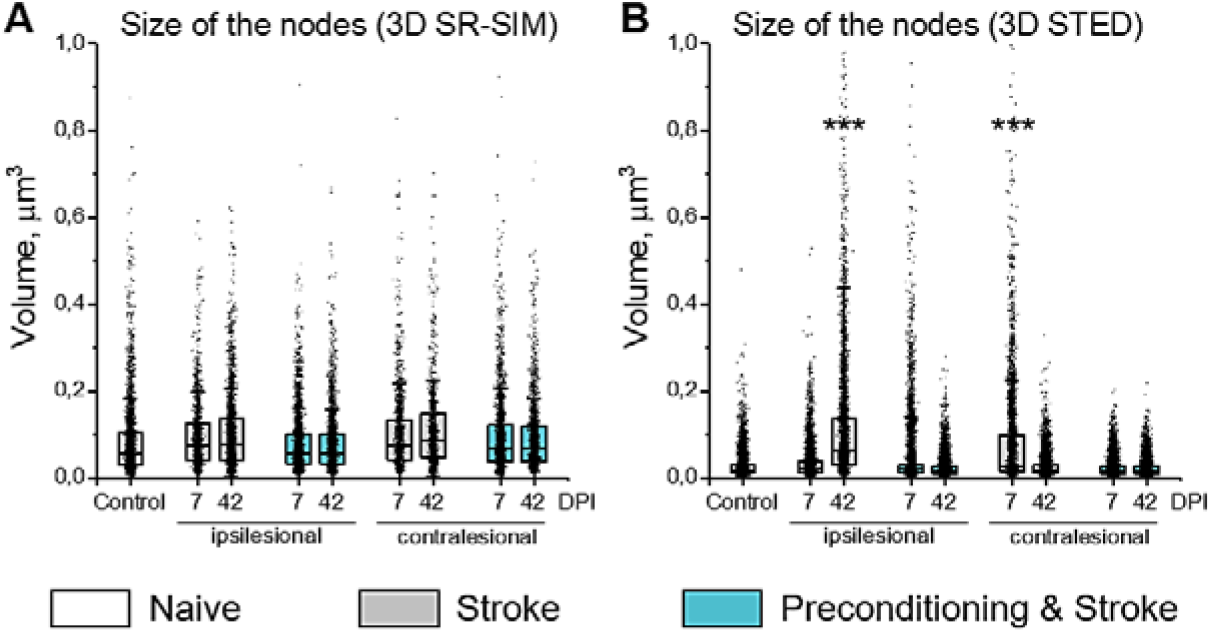
Size of PNN nodes post stroke. The size of PNN nodes was measured with superresolution (**A**) SR-SIM and (**B**) 3D STED microscopy as the volume of ellipsoids. Graphs are box plots with data as dots, means as squares, medians as lines, interquartile ranges as boxes and whiskers showing SD. Asterisks denote significant differences with the control group, as indicated by two-way ANOVA and t-tests (***p < 0.001), n=5. DPI, days post ischemia.

**Figure S5.**
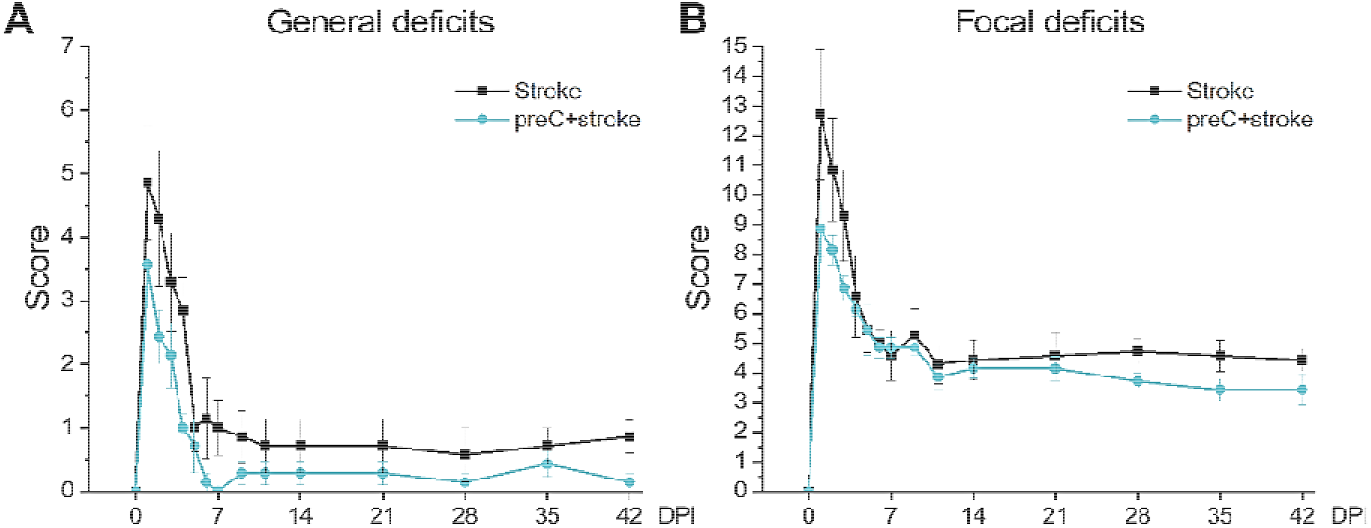
Neurological deficits post stroke. (**A**) Clark’s general deficits scoring. (**B**) Clark’s focal deficits scoring. Data are mean±s.e.m. n = 7. DPI, days post ischemia.

